# FluOpti: an open-source system for programmable whole-plate optogenetics and multi-fluorescence imaging

**DOI:** 10.64898/2026.10.12.758240

**Authors:** Isaac Nuñez, Tamara Matute, Alejandro Aravena, Wladimir Araya, Daniel Nuñez, Severine Cazaux, Valentina Ferrando, Patricio Obregón, Francisco J. Quero, Gustavo Düring, Timothy Rudge, Fernan Federici

## Abstract

Bacteria are able to self-organize into complex spatial communities, such as colonies, biofilms and swarms. Probing cells and gene networks with dynamic stimuli while preserving their spatial organization is therefore essential for understanding how bacterial communities organize across spatial and temporal scales. Optogenetics offers a powerful approach for such a task given the precise control of stimulus delivery without the perturbations normally associated with the addition of liquid inducers. However, most optogenetics devices have been developed to work with liquid cell cultures or to deliver a spatial pattern without the concurrent recording of gene responses and colony growth. Here, we describe FluOpti, an open source, low-cost device for controlling gene expression, through programmable light inputs, and recording multi-fluorescence responses and colony growth across whole petri dishes. White light LEDs are used for brightfield imaging while green and red LEDs are used as optogenetic inputs to activate and inactivate target genes via the CcaS/R two component system in *E. coli*. Low cost blue and orange acrylic are used as excitation and emission filters for fluorescence imaging, respectively. We developed custom software to analyze colony growth and optogenetic responses from 72-hour time-lapse experiments. A mathematical model was developed to predict gene expression levels in response to green-to-red light ratio stimuli. We use FluOpti to reveal a spatially structured response to optogenetic stimuli, with gene activation rates differing in cells located at the center and edge of colonies. to validate its use at a whole petri dish scale, FluOpti was used to depict how Voronoi territories can act as predictors of colony size, in congruence with previous studies. Together, these results demonstrate the utility of the device for studying gene networks at multiple spatio-temporal scales, from individual colonies to the whole petri dish. Performing optogenetics studies in whole petri dishes while preserving spatial arrangements could offer valuable insights into the spatial biology of gene networks in microbial populations.

**Graphical Abstract:** 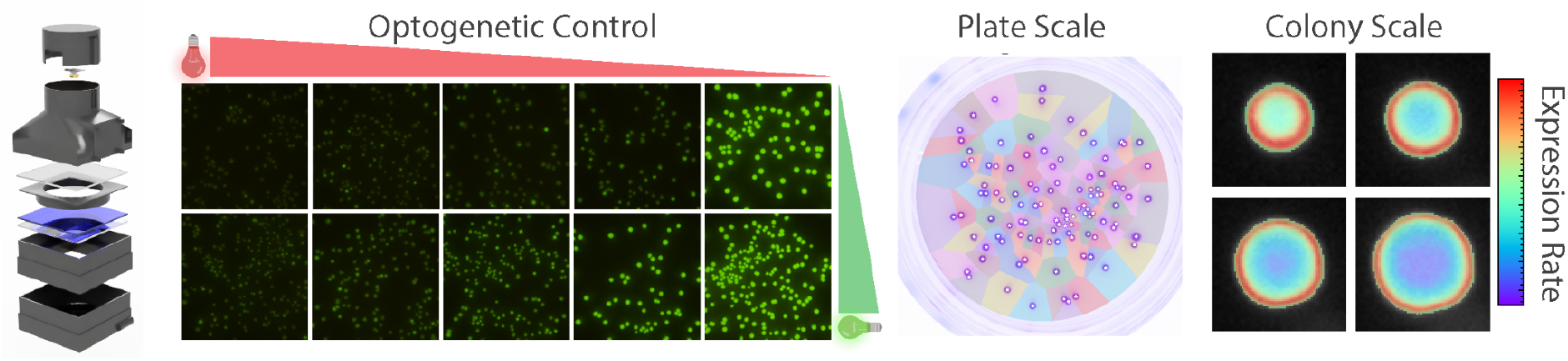

## Introduction

Bacterial populations self-organize into complex multicellular structures when grown on solid media, ranging from colonies to biofilms and swarms. Growth on solid surfaces enables bacterial ensembles to access unique ecological dynamics and spatial organization not present in well mixed liquid cultures [1–4]. Even simple genetic circuits have been shown to exhibit complex behavior when embedded in spatially distributed bacterial colonies [5]. Understanding how these multiscale organizations emerge requires experimental designs able to interrogate the spatiotemporal behaviour of gene networks and cells within their intact spatial configuration. Applying dynamical stimuli to gene networks while preserving self-organized spatial interactions on solid media can complement synthetic biology approaches studying spatial behaviour at a colony [5–11], inter-colony [12–14] and whole plate scales [15,16]. While chemically-induced transcription systems, such as IPTG/LacI and arabinose/AraC, have been instrumental for gene network studies due to their ON/OFF high dynamic range and reliability (e.g. [17]), their use as dynamical stimuli on solid media is challenging.

Optogenetics, the use of light as a controller of genetically-encoded photosensors has emerged as a powerful tool for precise spatio-temporal regulation of gene expression in bacteria [18–24], yeast [25–27] and mammalian cells [28–32]. This progress in light-responsive genetic systems, ranging from transcriptional activators [33] to recombinases [34], has been accompanied by the development of specialized hardware to control light intensity in a time-varying manner [18]. The use of programmable electronics for optogenetic devices have led to significant progress in automated feedback control of biological processes [31,35–39] as well as a better characterization of optogenetic systems (e.g. [40]). Most of these devices, however, have been designed for optogenetic control of cells in liquid cultures, e.g. turbidostats [35,38,39], microfluidic chips [41,42], culture tubes [43,44] and multiwell plates [37,45–52]. However, the application of optogenetics to intact bacterial colonies on solid media has been less explored. While specialized equipment has been developed to deliver patterned light inputs from projectors or photomasks with remarkable resolution [53–59], these devices have not been designed to track growth and fluorescence across individual colonies on a petri dish. Therefore, establishing a causal relationship between input light and genetic responses from time lapse experiments remains challenging.

Here, we describe FluOpti, an open source, low cost and compact instrument to perform high-throughput optogenetics, colony growth measurements and multi-fluorescence imaging of bacteria growing on solid media. FluOpti combines capabilities of the FluoPi Raspberry Pi-based device for multi-fluorescence imaging from whole plates [60] and optogenetic LED illumination originally described for optogenetic control of the CcaS/R system [33,52,61] in liquid culture [35,43].

## Results

### 3D printed modular holding of optogenetic and imaging systems

FluOpti delivers programmable regimes of red and green light for optogenetic control of the CcaS/R system in bacteria, blue light for fluorescence readout and white light for brightfield imaging (Figure 1a, b, c). This device is composed of 3D printed modules that house (from bottom to top): a fan, blue illumination board, diffuser, blue filter, bottom ITO glass, petri dish holder, white illumination ring, top ITO glass, red/green optogenetic illumination, orange emission filter and Raspberry Pi camera (Figure 1a)(Figure S1). This conic design is inspired by an open source design used for transmitted white light imaging of whole plates [62] that provides perfect sealing for complete darkness in the absence of illumination. An openSCAD source code is provided for the whole design, allowing adjustments such as the height of cameras with different focal lengths or the distance between filters and illumination board for an even distribution of light (according to light angles of the LEDs). In addition, we developed 3D printed add-on modules for the FluoPi acrylic device [60], enabling an upgrade and recycling of this existing hardware (Figure S2).

**Figure 1:**
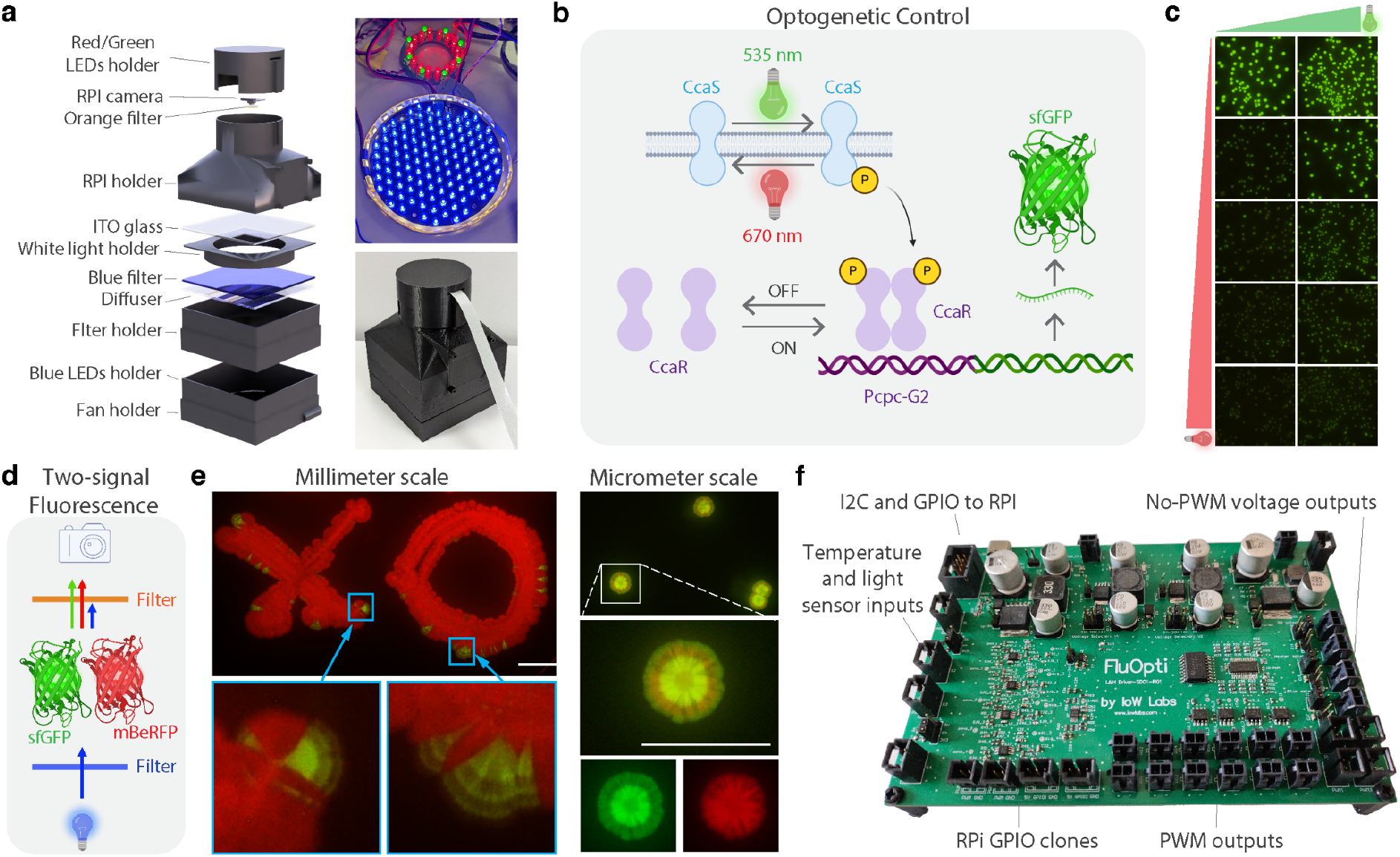
Open source hardware for optogenetic stimulation and multi-fluorescence imaging of the CcaS/R system. a) Schematic of FluOpti Modular Design (left) and image of 3D printed unit (lower right), showing the blue illumination base (upper right). A Raspberry Pi camera is used for programmable image acquisition. b) Schematic diagram of the CcaS/R optogenetic system that switches ON and OFF target gene expression in response to green and red light exposure, respectively. The CcaS is autophosphorylated upon 535 nm green light exposure, activating CcaR dimerization and activation of downstream genes under the cpcG2 promoter, sfGFP in this case. c) Whole plate imaging of colony responses to optogenetic signals of different green and red light intensity (from top to bottom: %0, %25, %50, %75, %100 of red light; from left to right: %0, %100 of green light). Raw images without editing. d) Left, schematic diagram of the arrangement of acrylic filters and illumination for multi-fluorescence imaging from a single illumination source. A blue filter is used as an excitation filter for the 470 nm LED-based illumination. Long Stokes shift fluorescent proteins, such as mBeRFP, are used concomitantly with sfGFP for red and green fluorescence. An orange acrylic is used as a long-pass emission filter to cut off wavelengths shorter than 500 nm (e.g. blue excitation light). e) Left, representative image of mBeRFP-expressing bacteria mixed with sfGFP-expressing optogenetic strain showing responses to 12h blocks of red and green light stimulation. Image levels linearly increased for clarity (original file available in the Zenodo repository). Scale bar = 10 mm. Right, concentric red/green states created by a regime of alternating 24-hour blocks of red, green, red and green light, respectively. Left, raw images with no editing. Scale bar=5 mm. f) FluOpti printed circuit board (PCB).

### Optogenetic system

We employed the CcaS/R two-component system [33], which responds to green and red light to activate and repress target genes of interest, respectively (Figure 1b, c). The CcaS/R system is a well characterized two-component system from *Synechocystis* PCC6803 that acts as a signal transduction pathway controlling target gene expression from the cpcG2 promoter upon green and red light exposure [61,63,64]. This two-component system acts via a sensor histidine kinase (CcaS) in the membrane that auto-phosphorylates upon green light exposure, transferring this phosphoryl group to response regulator (CcaR), which in its phosphorylated form binds to the cpcG2 promoter to activate transcription. CcaS is able to absorb light via a covalently ligated phycocyanobilin (PCB) chromophore. The engineered CcaS/R system for *E. coli* consists of two plasmids: pSR58.6 and pNO286.3, which produce all these elements [33]. CcaS is produced, from a J23106 constitutive promoter, as a green-light-sensitive inactive state, termed Pg, which switches to a red-light-sensitive active state (Pr) in response to green light [63]. Red light switches CcaS Pr to the inactive Pg state. CcaR is produced from a J23100 constitutive promoter. This photochromic system exhibits high dynamic range and fast kinetics [65].

Using a red- and green-light optogenetic system, instead of the more common blue-light-responsive circuits (e.g. [37,66–68]), frees the blue wavelength range for imaging fluorophores excited in this spectrum, such as GFP [60] (Video S1). To obtain a second fluorescent signal from the same blue light excitation, we employed a long Stokes shift fluorescent protein called mBeRFP [69], which emits in the red wavelength range. mBeRFP and sfGFP can be imaged simultaneously with a long pass filter cutting at around 500 nm (Figure 1d). This combination enables two fluorescent signals from a single excitation source (Figure 1e), avoiding the use of moving filter wheels that can introduce mechanical vibration in such a lightweight device [60]. This design is based on a tested configuration that successfully delivers high signal-to-background fluorescence for green, orange and red fluorescent proteins [60].

### FluOpti printed circuit board (PCB) integrating growth, optogenetic and imaging control

We created a printed circuit board (PCB) designed to integrate modules for temperature regulation, dynamic red/green light input programming, fluorescence microscopy and white-light imaging (Figure 1f)(Figure S3). The board contains a PWM generator at its core. The signals are fed to the drivers responsible for controlling output levels, e.g. for LEDs and heaters. The board features two types of onboard drivers, delivering four high power and eight low power channels, in order to accommodate different loads and consumption requirements separately. Depending on the configuration used (series or parallel, or some combination of these) the LEDs can have different operating points, including large loads (high voltages with high currents) that are difficult to handle by conventional drivers.

### Optogenetic illumination system

The optogenetic control of the CcaS/R two component system is performed with 520 nm 5mm and 660 nm Deep-Red 5mm LEDs. These LEDs are mounted on a custom-made circuit designed to be independently programmed, which sits as a ring surrounding the camera (Figure S4). We have developed two versions of this red/green LED board: one that can be manufactured in-house using low-cost CNC routers (Figure S4a) and another that can be produced by a commercial manufacturer as a printed circuit board (PCB) (Figure S4b). A 3D printed case holds the ring and a diffuser filter in place (Figure S4c).

In order to register colony growth, a lateral petri dish illumination was added for bright field imaging from a white LED strip. This module consists of a 3D printed ring that holds a white light LED strip (Forward current: 20mA @ 12V), surrounding the plate (Figure S5a) (Video S2).

### Fluorescence illumination system

Blue illumination for multifluorescence imaging is delivered by a LED board designed to display 115 LEDs emitting at 470 nm (Figure S5b). A diffuser sits underneath the 3 mm–thick blue acrylic sheet to produce an even distribution of the blue light (Figure S5c). A 126 × 126 mm laser cut blue acrylic serves as the excitation filter (Figure S5d), while a 3 mm–thick orange acrylic, laser-cut to a 1 inch diameter, is used as the emission filter (Figure S5e). This filter can be positioned in front of three Raspberry Pi camera models (HQ, M12 5MP and 8MP) using 3D printed adapters that also hold the camera (Figure S5f). The design is optimized for the version 2 of the Raspberry Pi camera, providing 3D printed tools to adjust its focal distance by lens rotation. The model is also compatible with the high resolution Raspberry Pi camera module 3, with digital control of focus and autofocus (Figure S5g). To evaluate excitation uniformity, an array of spatially distributed 2 µL samples containing 0.01 mM fluorescein was imaged across the full surface of the Petri dish. This concentration is within the range of fluorescence intensity obtained for sfGFP in our experiments (data not shown). We observed a mean pixel intensity of 106.14 with a standard deviation of 3.50, obtaining a coefficient of variation of 0.033, indicating a spatially uniform distribution of fluorescence intensities (Figure S5h).

### Temperature control

The FluOpti board incorporates an analog to digital converter, controlled with an I2C protocol, for reading 4 analog temperature sensors (NTC), and 2 resistive heats, allowing up to two automatic temperature controls to be implemented (Figure S6). These elements are used for PID temperature control via a transparent conductive glass coated with Indium Tin Oxide (ITO) (<7 ohm/sq, 100×100×1.1 mm), sitting below and above the petri dishes (Figure S6b).

### User interface

To integrate optogenetic and fluorescence imaging capabilities, we developed a user interface that facilitates the elaboration of optogenetic and imaging routines. An API was developed in Python to implement low-level control of the modules available on the FluOpti board. It allows the configuration of power of each output channel and data reading from the sensors via the ADC available on the board. From these simple functions, complex protocols can be implemented, including camera setup, temperature control loops and LED intensities (Figure S7).

### Google Colab analysis modules

We developed a pipeline in Google Colab notebooks for image processing and data analysis (Suppl. Note 1; See https://github.com/LabTecLibres/FluOpti). The procedure starts by extracting information regarding pixel to mm conversion using the known dimensions of the petri dish (Figure 2a). The timelapse images are first subjected to a Gaussian Filtering procedure, which users can adjust in an interactive frame (Figure S8). Additionally, this code can identify and correct minimal frame displacements introduced by accidental movements of the instrument or vibrations during the time lapse execution (Suppl. Note 1). Colonies are automatically identified as Regions of interest (ROIs) in the last frame of the time-lapse. These ROIs are then used to identify the same colonies across time frames. The procedure allows users to manually select or deselect colonies according to their requirements (e.g., analyzing only colonies that have not collided or filter out artifacts that may be detected in the image) (Figure 2b) (see Supplementary Note 1 for a full description of the pipeline). Colony edges are detected in brightfield images by a parametrized function that allows threshold adjustments (Figure 2c), and individual colony growth is calculated across time frames (Figure 2d, e) (Video S3, Video S4). Automated colony selection is comparable to manual selection for plates populated with less than 200 colonies (Figure 2f). This density range is appropriate for individual colony studies as well as whole plate analyses, such as testing the effect of the Voronoi area on colony size variation, which has been found to be a good predictor of colony size variation [70,71](Figure 2g,h)(Table S1).

**Figure 2:**
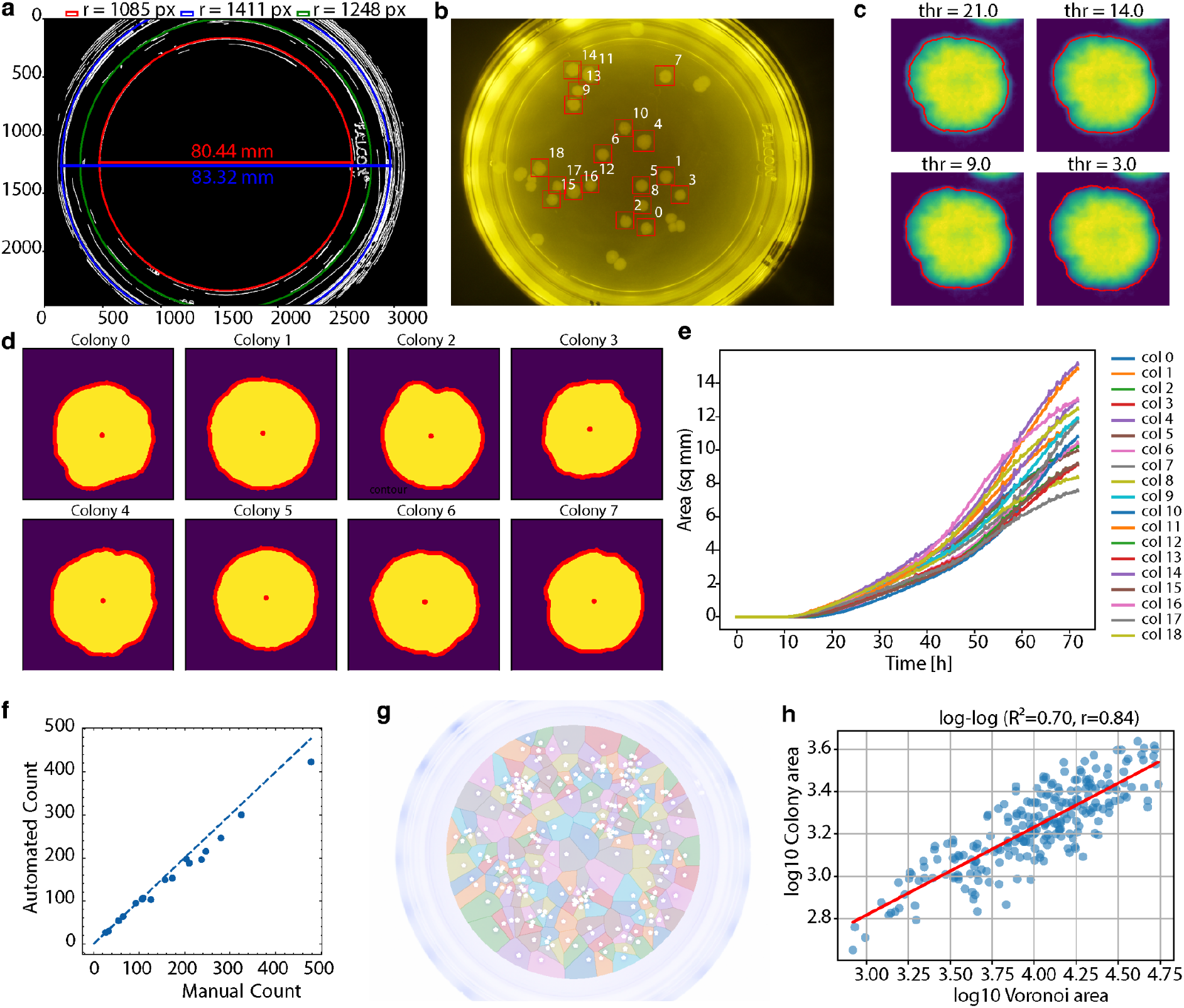
Colony tracking and analysis. a) Pixel to mm conversion using the known width of petri dishes. b) Regions of interest defined for each colony, considering distance to other colonies. c) Different thresholding for brightfield image of colony #9. d) Areas calculated colonies 0 to 8 as an example (Threshold=21, t=72h). e) Changes in colony area per time in a 72h time lapse experiment (threshold = 7). f) Comparison of colony counts obtained using automated image analysis versus manual quantification. g) Voronoi segmentation of agar territory available to each colony. h) Scatter plot showing the relationship between Voronoi cell area and colony size. The correlation coefficient and statistical significance (p) are indicated (p-value << 0.05).

### Validation

To assess the performance of FluOpti in high-throughput optogenetic experiments, we conducted a series of time lapses consisting of different routines of optogenetic stimulation. First, we applied 24-hour blocks of red and green illumination for 72 hours (Figure 3a). As the proliferation of cells is higher at the edge of colonies, giving rise to linear colony growth after 16h (Figure 2e), induction-repression cycles revealed different responses in cells positioned at the edge versus center of the colony. The fluorescence profile in the center of the colony induced by 24h of green light remains almost intact after 10h of red light repression (Figure 3a,b), indicating that transcriptional shutdown is not enough to reduce GFP levels in this region of the colony. In the absence of degradation tags, GFP kinetics is predominantly regulated by induction (and dilution) and not by protein turnover. To study whether cells at the center of the colony are still able to respond to induction stimuli, we implemented a protocol consisting of 36h of red light followed by 36h of green illumination and quantified the rate of optogenetic responses within colonies (Figure 3c). All cells of the colony responded immediately to the green stimulation, to change later to a ring of higher expression rates at the edges. Experiments with continuous green light stimulation also revealed the presence of this edge ring and the transition from whole colony to edge expression that coincides with the transition of exponential to linear colony growth (Figure 3d)(Figure S9). As colony edges are defined from brightfield images, non-fluorescent cells at the edge would serve to reveal protein dilution due to cell proliferation. Consistent with this, optogenetically induced GFP levels are diluted by cell proliferation at the colony edge, forming concentric ring patterns when the colony is subjected to cycles of 12h red light after 4h of green light induction (Figure 3e).

**Figure 3:**
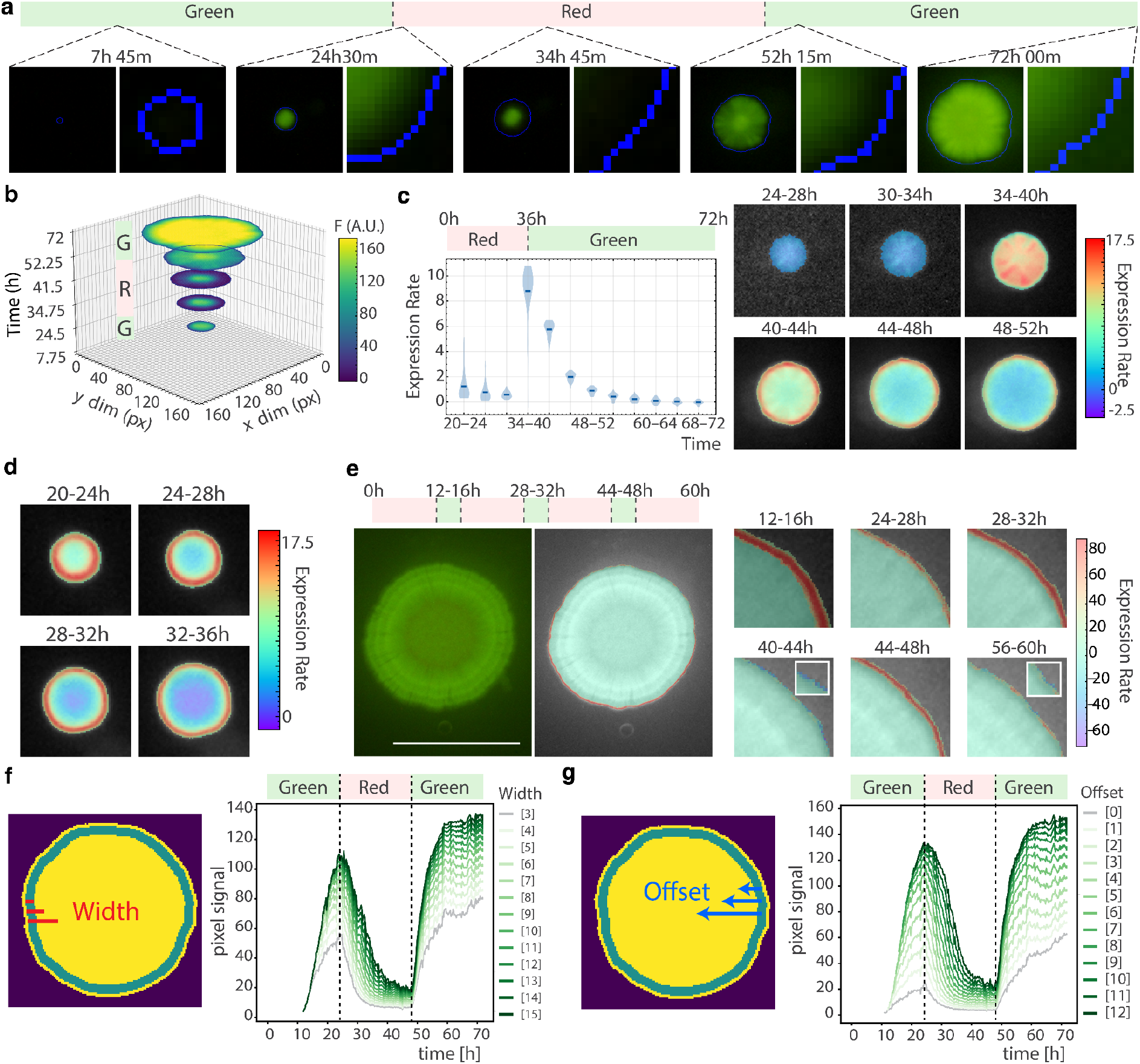
Optogenetic performance. a) Representative colony from a 72h time-lapse experiment consisting of 24h blocks of green, red, and green illumination. The colony edge is defined in blue from the brightfield channel, accounting for all cells in a colony regardless of their fluorescent signal. A zoomed in area is shown in the right panels for each time point. b) Fluorescence signal with respect to colony size at the time points shown in (a). R and G refers to red and green light optogenetic stimulation, respectively. c) Left, mean GFP expression rates in colonies subjected to green light stimulation after 36h of red light repression (Solid blue line: Mean of the whole colony across the 4 hour interval range). Right, representative colony showing expression rate distribution in response to green light stimulation after 36h of red light repression. Heatmap bar: fluorescence per hour.. d) Rates of GFP expression under constant green light stimulation. Rates were calculated using 2 hour blocks (8 data points per block) by taking the difference between the fluorescence images of the ROI at each point and averaging them across the block. Heatmap: fluorescence as pixel intensity per hour. e) Rates of GFP expression in response to an initial step of 12hours of red stimulation followed by 3 cycles of 4h of green light and 12h of red light. Scale bar= 10 mm. f) Representative analysis of intensity values at different ROI widths (3, 4, 5, 6, 7, 8, 9, 10, 11, 12, 13, 14, 15 pixels) with a fixed offset of 1 pixel. The top schematic diagram shows a width (red) of 0.20 and offset of 0.05. Red and green steps represent light input conditions, i.e. red and green light. g) Representative analysis of intensity values at different ROI offsets (0, 1, 2, 3, 4, 5, 6, 7, 8, 9, 10, 11, 12 pixels) with a fixed border width of 1 pixel. The top schematic diagram shows an offset (blue) parameter of 0.05 and width of 0.20. Red and green steps represent light input conditions, i.e. red and green light.

To characterize the optogenetic responses in the edge better, we define ROIs that capture exclusively this cohort of cells. A ring of pixels was defined at different positions from the border to the center of the colony (offset values) and containing different numbers of pixels at both sides of the ROI (width values) (Figure 3f, g). Once defined, the position of this ROI is updated at each time frame, while its width remains fixed. The fluorescence signal increases with the ROI width because more pixels from the colony interior are included, as observed in colonies continuously stimulated with green light (Figure S10). The offset value serves as a trade-off parameter for dynamic measurements as intensity rises with pixels selected further into the center and away from the colony edge whereas rates of expression decreases: larger offsets capture higher signals from the colony center, whereas smaller offsets focus on cells proliferating at a constant rate near the colony edge (Figure 3f,g).

In order to characterize optogenetic responses in these edge cohorts, we created a mathematical model that describes fluorescence dynamics in response to light stimulus, based on previous work by [72] and [60] (see Supplementary note 2 for model details), assuming constant growth and dilution of protein and mRNA molecules. The model incorporates the chemical reactions that describe the activation and inactivation of the CcaR transcription factor, its binding to the target promoter, and the subsequent transcription and translation processes, leading to the production of superfolder GFP (sfGFP) protein. The CcaR transcription factor can exist in an inactive OFF state or an active ON state, with transitions between these states controlled by the intensity of red and green light, respectively. Based on the results described above, we fitted the model to data corresponding to the fluorescence dynamics of cells located at the edge of the colony, where cells actively proliferate. We use this ODE for the model fitting procedure to the experimental data, by numerically integrating the differential equation, and using Bayesian parameter estimation techniques for estimating the parameters from our experimental data (Supplementary note 2).

The model demonstrated strong predictive performance on datasets that were not used for parameter estimation, achieving an overall R^2^ of 0.94 (n = 28,800 single time point observations; RMSE = 11.9) (Figure 4a). Posterior parameter distributions and inferred parameter correlations from the Bayesian parameter estimation process are reported in Figure S11. The predictions accurately captured the fluorescence dynamics under a regime of 60% green light intensity, however, the predictions for the 100% red light regime without green light showed reduced accuracy (R^2^ = 0.29). This could be explained due to the fact that this condition represented extrapolation beyond the training data range, which included red light intensities up to 75% without green light or 100% red light plus green light, but not 100% red light illumination on its own. Further analysis showed that despite the low R^2^, the model predictions remained strongly correlated with our experimental data (r = 0.80). This added to the fact that the data showed high variability (CV = 84%) suggests that the reduced R^2^ found for this dataset would reflect a systematic positive bias (mean residual for the dataset = +3.3 A.U.), slightly overestimating the absolute fluorescence levels. This indicates that the proposed ODE model can correctly capture the quantitative temporal patterns within the training matrix. The model was compared to experimental data consisting of 24 h blocks of activation-repression cycles (Figure 4b). These routines induce sharp activation and deactivation of fluorescent signals similar to those already described in liquid cultures [43] (Figure S12).

**Figure 4:**
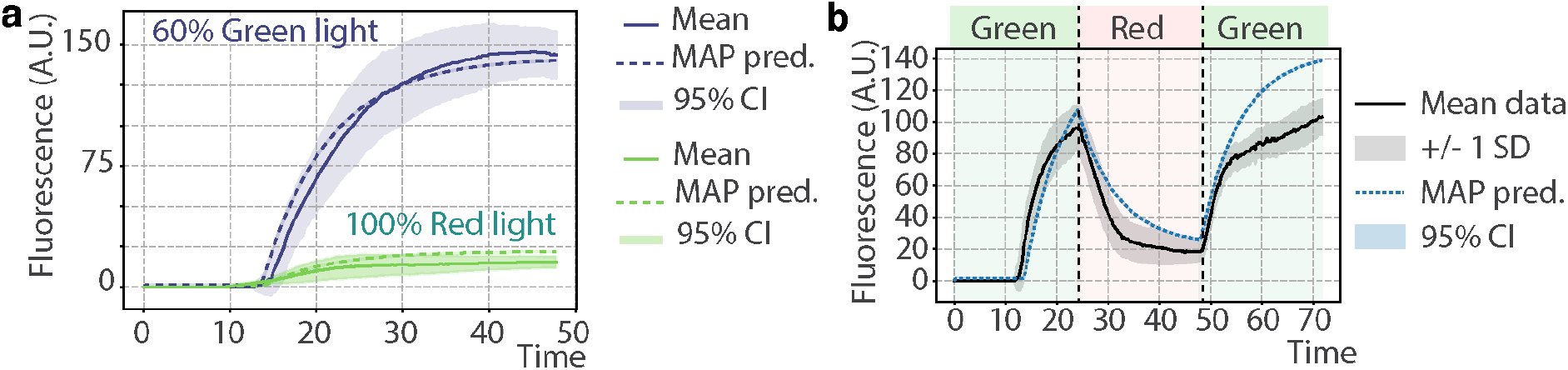
Mathematical model. a) Model prediction of optogenetic responses. Model predictions for data that was withheld from initial training. Dashed lines indicate the model’s prediction, continuous lines correspond to the mean of the experimental data. b) Model prediction of multiple regimes. Model prediction of a varying light regime, corresponding to three blocks of 24 hours of green, red and green light, respectively. Blue dashed line: Model prediction. Black continuous line: Mean of experimental data.

## Discussion

In this work, we describe FluOpti, an open-source device for high-throughput optogenetic control and multi-fluorescence imaging of microbial colonies. The main advantage of FluOpti is the possibility of studying gene regulatory dynamics across large spatio-temporal scales that emerge from self-organization of bacterial cultures on solid media. Applying dynamical stimuli while maintaining microbial organization intact is difficult with other approaches such as chemical inducers because it causes physical perturbations to the cells and the substrate as well as an heterogeneous accumulation and degradation of the inducer [73]. While lab-on-a-chip microfluidic methods offer a suitable approach for the dynamical characterization of gene networks (e.g. [74]), the required infrastructure for chip design, fabrication and operation remains inaccessible to many researchers in low- and middle-income countries. FluOpti offers a cost-effective alternative by leveraging microbial optogenetic systems [33,52,61,75] and recent developments in open source hardware for programmable red/green stimulation of liquid cultures [35,43], adapted here to enable optogenetics on solid media.

The advent of cheap microcontrollers and off-the-shelf electronics combined to 3D printing and laser cutting technology has enabled the development of a wide range of open scientific hardware (OSH), including microscopes, fluorescence imaging devices, PCR thermocyclers and plate readers [76–80]. The online sharing of design files and project documentation [81–84] under open licenses grants freedom to access, use, modify and distribute source code, design files and derived works, enabling easy reproduction by other users. Thus, the development of OSH devices not only result in cost savings of up to 94% compared to their commercial counterpart [85] but also give rise to instruments not found commercially, such as a whole-plate fluorescence imaging with optogenetic control. The use of open shareable design also gives rise to more collaborative practices and crowdsourced improvement of these research tools [86]. Here, we combine low-cost illumination, open source hardware for fluorescence imaging and a custom-designed board interacting with a Raspberry Pi computer. We designed a PCB that integrates the optogenetics hardware to a low-cost setup for multifluorescent imaging in order to have a standalone system. This approach circumvents the difficulties of retrofitting commercial microscopes, which is challenging due to inaccessibility to the proprietary software for hardware integration and customization. These microscopes are also subjected to high demand in university core facilities, which restricts opportunities for customization and long experiments. FluOpti applies whole plate multifluorescence imaging with a Raspberry Pi camera and custom software for segmentation and analysis. This approach also enables high throughput colony time lapse experiments that would otherwise require a mechanical stage for X-Y position tracking.

Thus, FluOpti allows long optogenetic procedures of hundreds of colonies at once, maintaining identical conditions that facilitate high-throughput experiments at a colony scale. This approach revealed how optogenetic activation is modulated by colony organization. Peripheral cells exhibit higher activation rates than cells located at the colony center. At the edge of the colony, constant access to nutrients and minimal mechanical constraints support active cell proliferation that dilutes GFP, leading to undetectable levels of fluorescence in response to red light treatment. Conversely, at the colony center, the lack of degradation tags in GFP leads to an irreversible accumulation of the signal. Given that optogenetic repression with red light does not produce equally rapid phenotypic reversal in all cells of the colony, a mathematical model was fitted to data obtained from the cohort of cells proliferating at the edge. The interplay between protein stability and the position of cells within the colony underscore the relevance of interrogating gene networks while embedded in their native collective organization. This is key for identifying spatially structured phenomena and interpreting optogenetic patterns quantitatively, which could be crucial to understand pattern forming circuits in whole colonies [7]. Analysis at a plate scale also permits studying colony behaviour with respect to spatial distribution on plates, such as Voronoi territory size as a predictor of colony size [70,71].

FluOpti provides an accessible tool for spatiotemporally resolved gene network studies that could be instrumental for revealing emergent phenomena in a wide range of microbiology studies; for instance those focusing on collective cell responses [2,68,87,88], microbial ecology [2,89–91], spatial organization of metabolic activities [92], microbial spatial organization [93,94], evolutionary mechanisms [3,95], morphogenetic engineering [96] and pattern forming gene networks [7,8,96–98].

## Material and methods

### Hardware

All schematic, design files and bill of materials can be found in the following repository: https://github.com/LabTecLibres/FluOpti, which is constantly updated.

The optogenetic illumination is based on 520 nm (Kingbright WP7083ZGD/G) 5mm LEDs and 660 nm Deep-Red (LEDSupply L2-0-R5TH50-1) 5mm LEDs. The orange acrylic filter (High-gloss, UV absorbent, cut tolerance of +/- 1/8 in, transparent, thickness of 1/8 with thickness Tolerance of +/- 8%) was obtained online from inventables.com [99]. The blue acrylic (#2424) was sourced from E & T Plastics of Florida (Ft. Lauderdale, FL 33309). Tracing paper (“papel diamante” or “papel vegetal” in Latin America) was used as a diffuser filter. The ITO glass used for temperature control is a 100x100x1.1 mm ITO with a resistance of <7ohm/sq.

### Genetic constructs

Genetic constructs used for this study were provided by Jeff Tabor [REF], and include the pNO286-3 plasmid which expresses CcaS, ho1, and pcyA ([pNO286-3](https://benchling.com/s/seq-41hQr2lcy42Wa9WvcDg1?m=slm-KnCU5VOVbeKS0AsiEQKc)), and the pSR58.6 plasmid which expresses CcaR constitutively, and sfGFP under the PcpcG2-172 promoter ([pSR58.6](https://benchling.com/s/seq-AbxDCK2sb1711MuxQB5E?m=slm-ZKjgVzjd2YouKXCIEyQl)). For the concentric rings of sfGFP and BeRFP shown in figure 1, we used the pOIR34 plasmid ([pOIR34](https://benchling.com/s/seq-7vWk1pxyjY11POnztBdV?m=slm-AjqAuAAawXVABBPSI2ax)).

### Plate cultures

*E. coli* streaks of strain BW29655 co transformed with the pNO284 and pSR58 plasmids were prepared and grown overnight with a full red light regime (16-18 hrs growth under red light) on solid LB media (10g of tryptone, 5g of yeast extract, and 5g of NaCl dissolved in deionized water to a final volume of 1L) or M9-Glucose (1 x M9 salts supplemented with 2 mM MgSO_4_·7H_2_O, 0.1 mM CaCl_2_, 0.4% Glucose and 0.2% Casaminoacids; 5 x M9 salts consist of 64g of Na_2_HPO_4_·7H_2_O, 15g of KH_2_PO_4_, 2.5g of NaCl, and 5g of NH_4_Cl dissolved in deionized water to a final volume of 1L), either one supplemented with 1.5% w/v agar (and 1% for Fig. 1e). Colonies from these plates were selected randomly and grown on the corresponding liquid media overnight, and these cultures were diluted 1:100.000 or 1:1.000.000 depending on the initial size of the selected colony (1:100.000 for colonies up to 1.5 mm diameter; 1:1.000.000 for colonies with a diameter larger than 1.5 mm). These dilutions were then plated on fresh solid LB or M9-Glucose media supplemented with their corresponding antibiotics and placed in the FluOpti device for experimental data acquisition. Antibiotics were prepared as stock solutions of kanamycin (50 μg/μL), carbenicillin (100 μg/μL), chloramphenicol (10 or 25 μg/μL), or 100 μg/μL spectinomycin depending on the experiment.

For spatially patterned cocultures of Figure 1e, plates were prepared using the sfGFP-producing optogenetic strain BW29655 and a second *E. coli* BW29655 strain carrying pNO286-3 and the uLoop plasmid pSB-even assembled with L1-mBeRFP, which provides the red fluorescent reporter. Colonies from each strain were collected separately and resuspended in 50 µL of sterile Milli-Q water. These suspensions were mixed to prepare “red and green” ratios of 10:1 and 20:1. The mixtures were inoculated onto M9-glucose plates containing 1.0% (w/v) agar, supplemented with 100μg/mL spectinomycin and 25 μg/mL chloramphenicol. Cross and circular shape patterns were streaked using an inoculation loop and arranged in a 2 x 2 layout and then subjected to the standard 72 hour FluOpti program where plates were imaged every 15 minutes. For spot assays, 1 µL droplets of single or mixed populations were precisely deposited onto solid media in a 3 x 3 grid configuration [3] (*See* Figure S13). Plates were allowed to dry and absorb before incubation. Parallel control plates were maintained under constant red light, both kept at standard incubations conditions.

The usual FluOpti illumination program consisted of an initial continuous 24 hours of red light exposure, followed by three cycles of 4 hours of green light and 12 hours of red light, for a total duration of 72 hours (Figure 1e). Protocol variations in Figure 3 included continuous or prolonged red or green illumination, shorter green light pulses of 20 minutes or reduced initial red light periods of 12 hours. Plates were placed inside FluOpti for the experimental data acquisition.

## Acknowledgment

We thank Jeff Tabor for helpful discussions and for providing plasmids.

We thank Thingiverse developer soswow for the 3D digital model of Raspberry Pi Camera Module v2.1 (https://www.thingiverse.com/thing:2376448/files) used in our 3D schematics.

## Author Contributions

FF, designed research;

FF, GD, TJR supervised research;

AA, WA, IN, TM, DN, VF performed research;

FJQ assisted in code implementation and RPI3 camera integration.

AA, IN, TM, and FF analyzed data;

FF, AA wrote the manuscript.

All authors reviewed the manuscript.

AA Alejandro Aravena

WA Wladimir Araya

IN Isaac Nuñez

TM Tamara Matute

VF Valentina Ferrando DN Daniel Núñez

FJQ: Francisco Javier Quero

GD: Gustavo Düring

TJR Timothy Rudge

FF Fernan Federici

## Funding

This work was supported by the National Agency for Research and Development (ANID) through the ANID Millennium Science Initiative Program (ICN17_022), Concurso de Investigación “Avanza UC 2026” Concurso de Investigación “Avanza UC 2026”, Fondo de Desarrollo Científico y Tecnológico FONDECYT Regular 1211218 and FONDECYT Regular 1241452 awarded to Fernan Federici. We thank the support of the project for Technological Centers of Excellence with Basal Financing ANID-Chile to the Cape Horn International Center–CHIC (ANID PIA/BASAL PFB210018) & Centro Ciencia & Vida, FB210008, Financiamiento Basal para Centros Científicos y Tecnológicos de Excelencia.

### IA declaration

During the preparation of this work the author(s) used Consensus for literature search and ChatGPT for grammar corrections and improving readability of the manuscript. Authors used GitHub Copilot to assist programming the Jupyter Notebooks. After using these tools, the author(s) reviewed and edited the content as needed and take(s) full responsibility for the content of the published article.

