## Supplementary Information for "FluOpti: an open-source system for programmable whole-plate optogenetics and multi-fluorescence imaging"

Supplementary Figures
Supplementary Table
Supplementary Videos

### Supplementary Figures

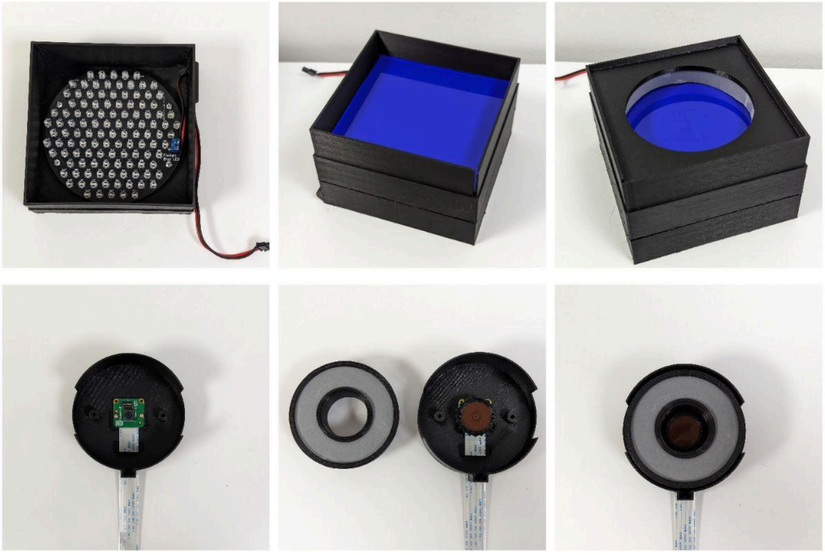

**Figure S1: FluOpti 3D printed case.** Top, Organization of base modules: Lighting base containing the PCB array of blue LEDs on top of a 12V fan (not shown), a blue filter on top of a diffuser (not shown) and a white illumination ring for brightfield imaging. Bottom, top part holding the camera, orange filter and the ring PCB holding the red and green LEDs, plus a diffuser for even optogenetic illumination.

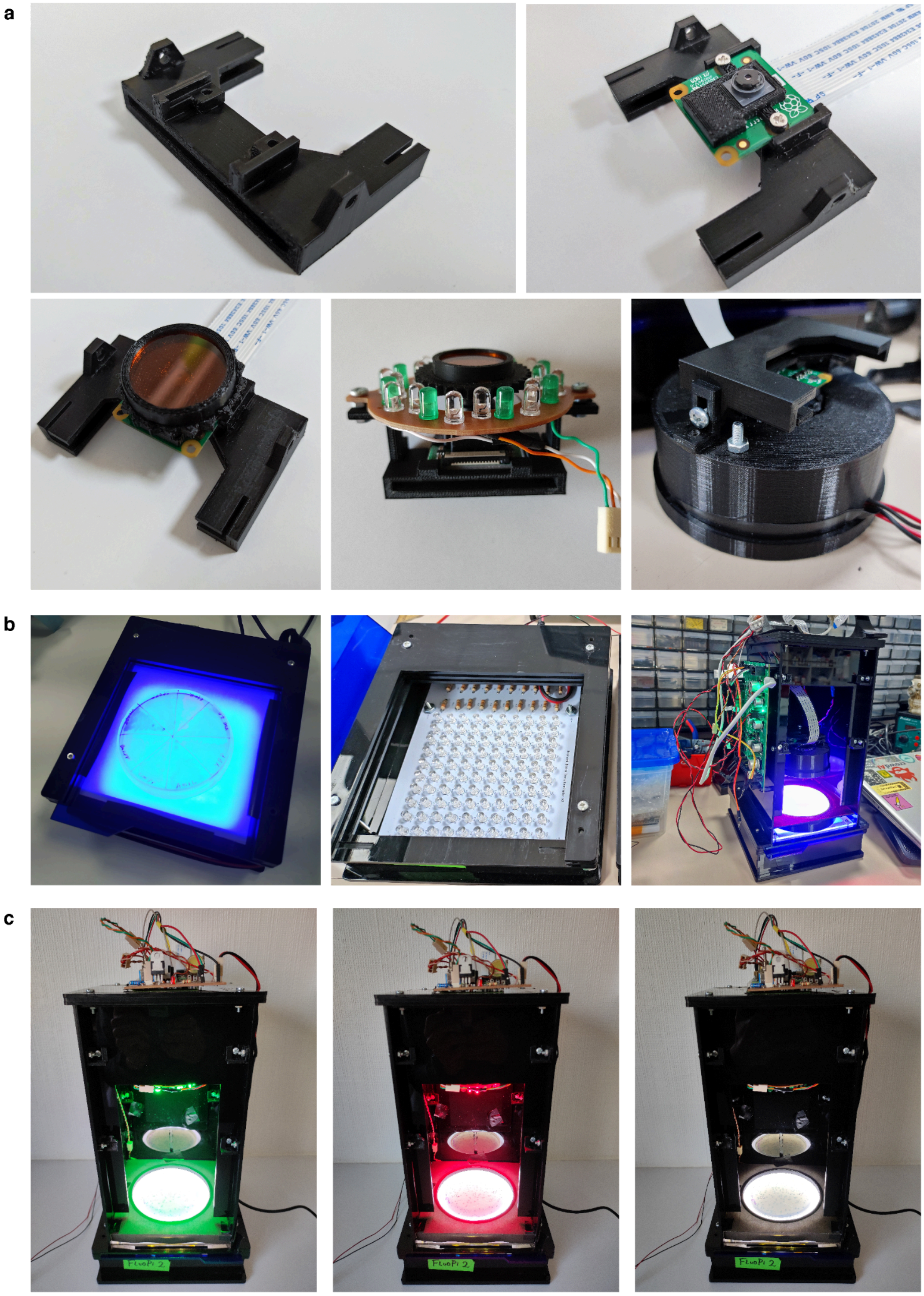


**Figure S2: FluoPi optogenetic upgrade.** a) Holder for camera and red-green LED circuit. b) The blue illumination is achieved with the same hardware described in [60]. c) Complete system showing an optogenetic routine and white illumination in sequence.

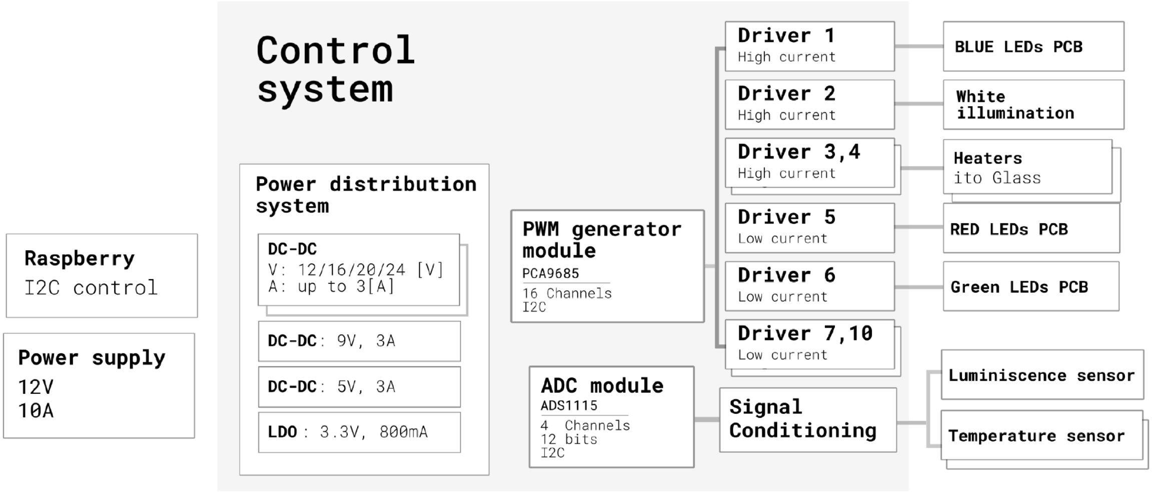


**Figure S3:** Schematic representation of FluOpti board control system and interaction with input/output modules. The main characteristics of the board are: 12V 10A power supply; 2 configurable output voltage sources for different operating loads; 2 channels for high consumption LEDs with selectable V and up to 2.5 A per channel; 2 channels for heaters (high consumption); 8 channels for low consumption LEDs with up to 500mA per channel; 2 PWM access signals from the Raspberry Pi board for general purpose use; up to 4 analog input channels to read temperature sensors; a 5V 2.5A output to power a raspberry; and system control by I2C. The board is compatible with different versions of Raspberry Pi. A set of five DC-DC converters generate a total of eight different voltages from a single 12V input supply. Two step-up DC-DC converters allow the generation of high voltages (12V, 16V, 20V and 24V) with the capacity to manage up to 3 A each. Each of these power rails can be selected to power the four output channel drivers, which can be used to control both high consumption LED configurations or heaters via PWM. Three step-down DC-DC converters generate voltages of 9V and 5V, each with a capacity of up to 3 amperes. The 5V channel is arranged to power a Raspberry 3b+ or Raspberry 4, which simplifies the number of connections necessary in the system, thus saving one power supply. Finally, a low-dropout (LDO) converter generates the 3.3V voltage necessary to power the digital components used. Once the operating or rail voltage has been selected, the PWM module drives these voltages through transistors. There are two types of drivers: low current and high current. The low current channels support up to 500mA and the high current channels allow controlling loads of up to 2.5A. Extra PWM signals are also available from the board for general purposes. Additionally, the system replicates two PWM signals coming from the raspberry in case of need. All digital components available on the board are controlled through the I2C protocol to simplify the number of cables and connections used.

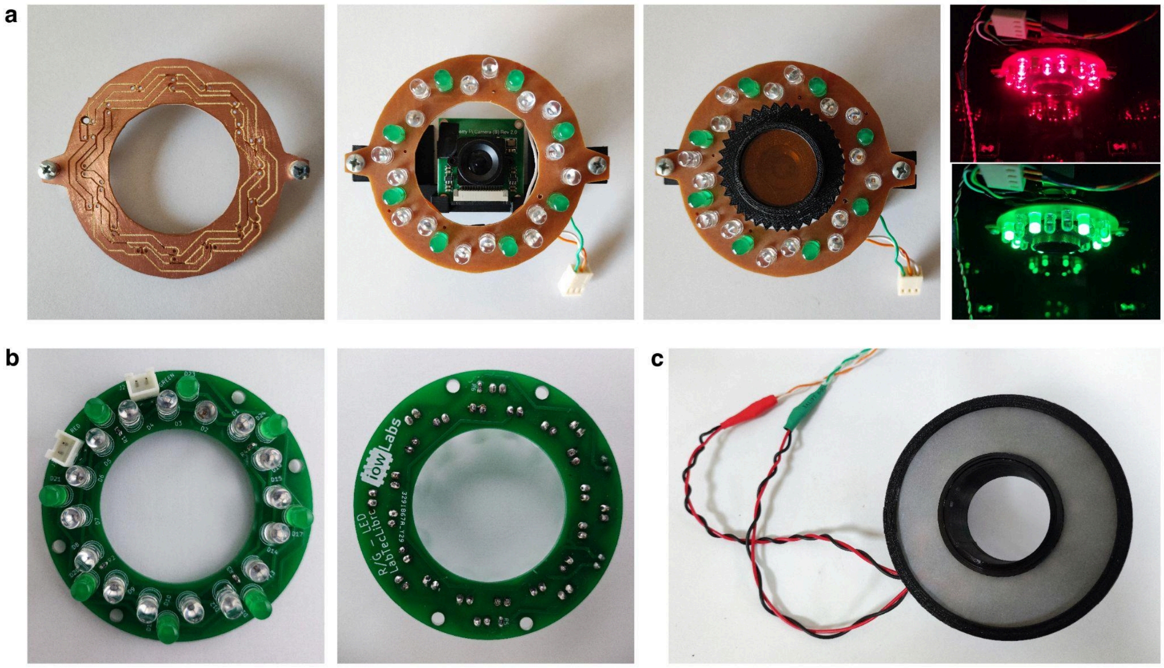

**Figure S4: Optogenetic module.** An ultra low cost optogenetic LED ring created with an open source CNC router (a) and a Printed circuit board (PCB) for the optogenetic LED module, custom-designed and produced by a commercial manufacturer (b), were created to accommodate the 520 nm Kingbright WP7083ZGD/G 5mm LEDs (Forward current: 20mA, at 3.2V. Peak forward current: 100mA, Max voltage forward: 4V) for green illumination and the 660 nm Deep-Red 5mm LEDs (Forward current: 20mA, at 2.2V. Peak forward current: 100mA, Max voltage forward: 2.8V) for red illumination. c) 3D printed case to hold a diffuser in front of the red and green LEDs for even illumination.

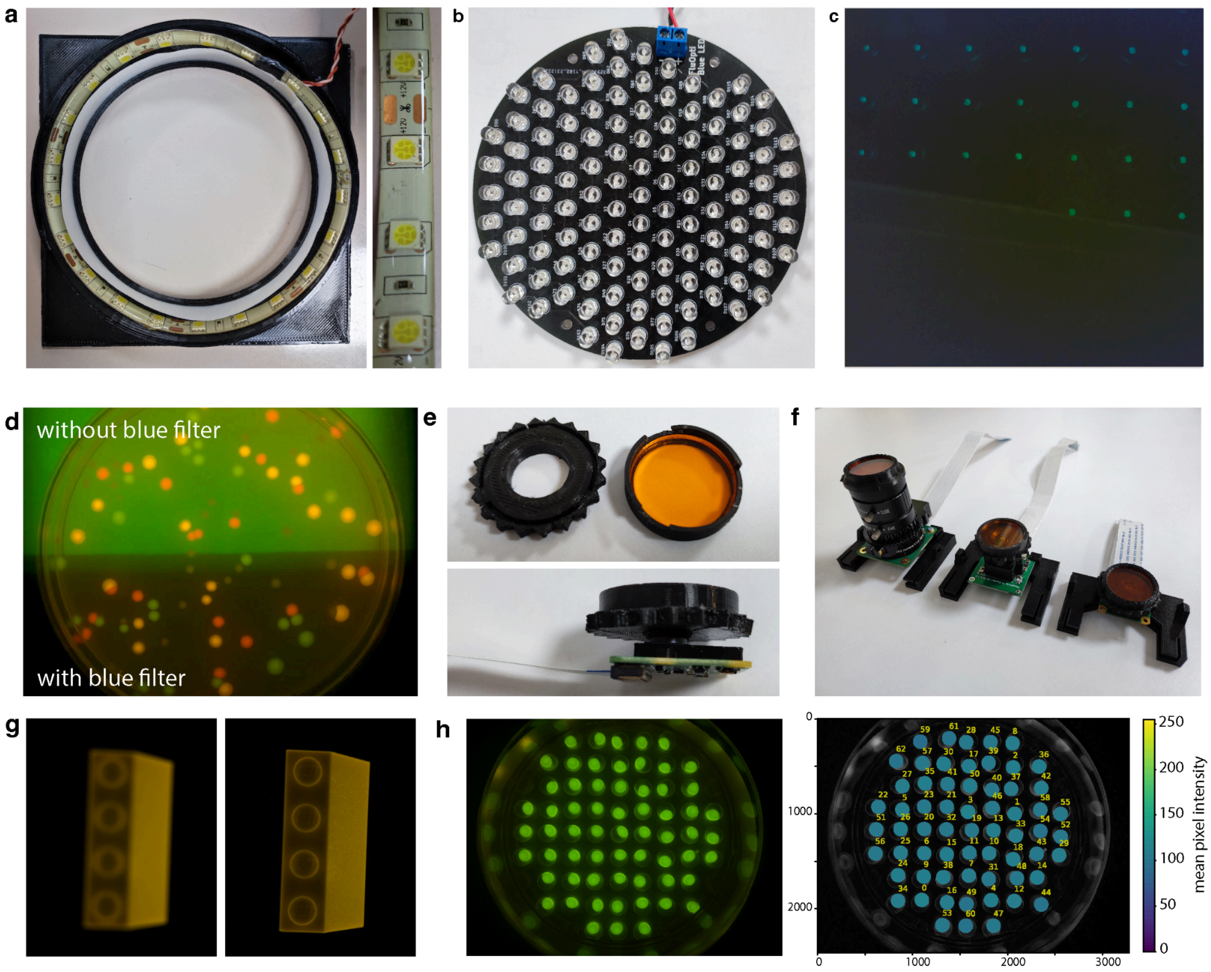


**Figure S5: Fluorescence and white light imaging module.** a) White light module housed in a 3D printed case. b) Custom-made PCB with mounted 470 nm blue LEDs (SuperBright, Forward current: 30mA at 3.3V. Peak forward current: 100 mA, Max voltage forward: 3.8 V). c) Diffuser moved half way over the LED array. d) Image of a plate sitting on top of diffuser and blue LEDs with (bottom) and without (top) the blue 3 mm acrylic used as the excitation filter. e) 3D printed holder and focus knob bearing a circle 3 mm thick orange acrylic used as an emission filter. f) Holder sets for different RPI cameras: HQ, 5MP adjustable focus with an 12-mm threaded objective lens (M12) and 8MP v2 RPI camera. g) Digital focus control of Raspberry Pi camera module 3. h) Fluorescence uniformity analysis from 2 µl samples of 0.01 mM of fluorescein (imaged at shutter speed of 25000 microsecond).

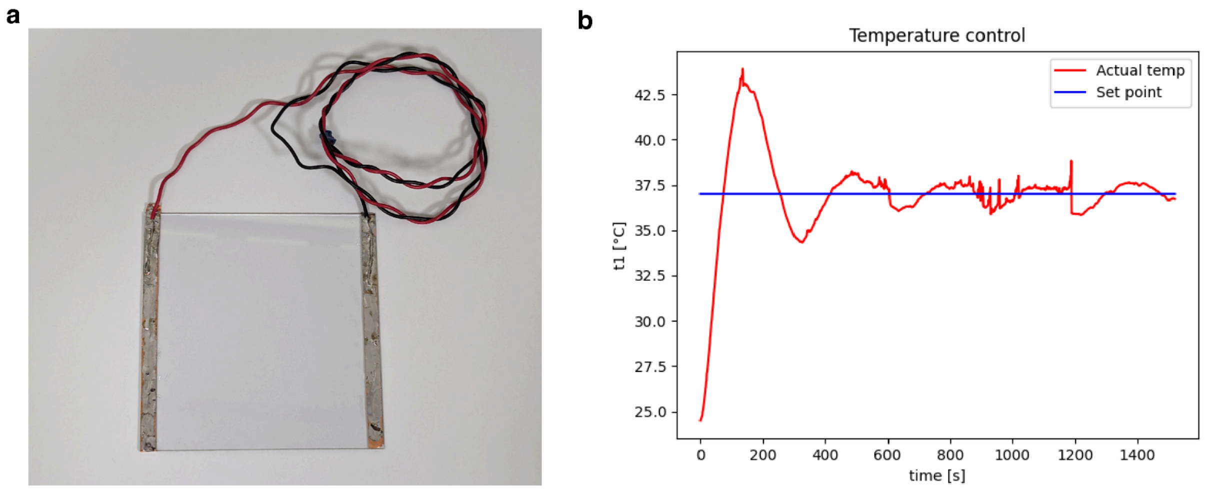


**Figure S6: Temperature control module.** a) Transparent conductive glass coated with Indium Tin Oxide (ITO) for temperature control. b) Plot showing 37°C output control.

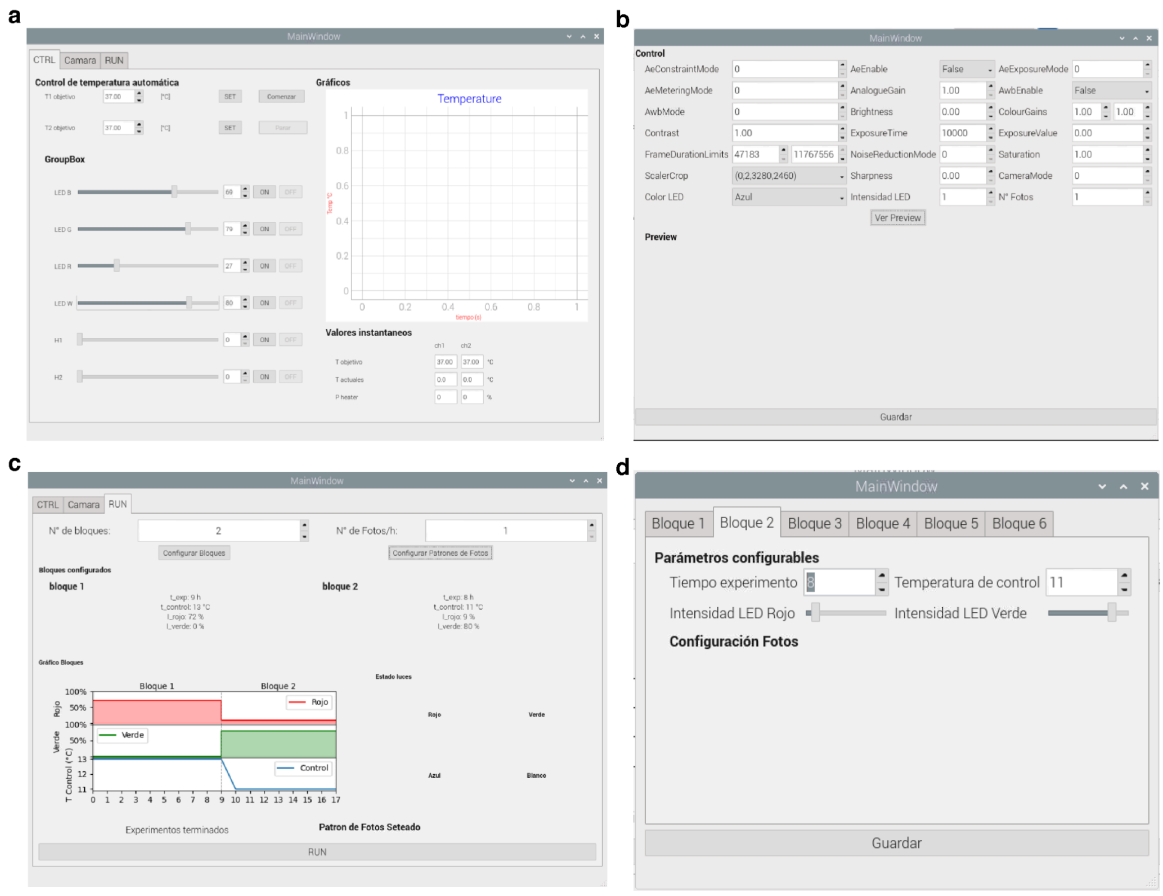


**Figure S7: GUI interface.** The GUI interface, designed using QT and pyqt5, allows the visual configuration of all the parameters of interest associated with FluOpti, such as LED intensity, heat Intensity, target temperature as well as the configuration for different types of image acquisition. a) Temperature and LED intensity control via GUI. b) Camera control parameters. c,d) Red/Green Routine setting via GUI.

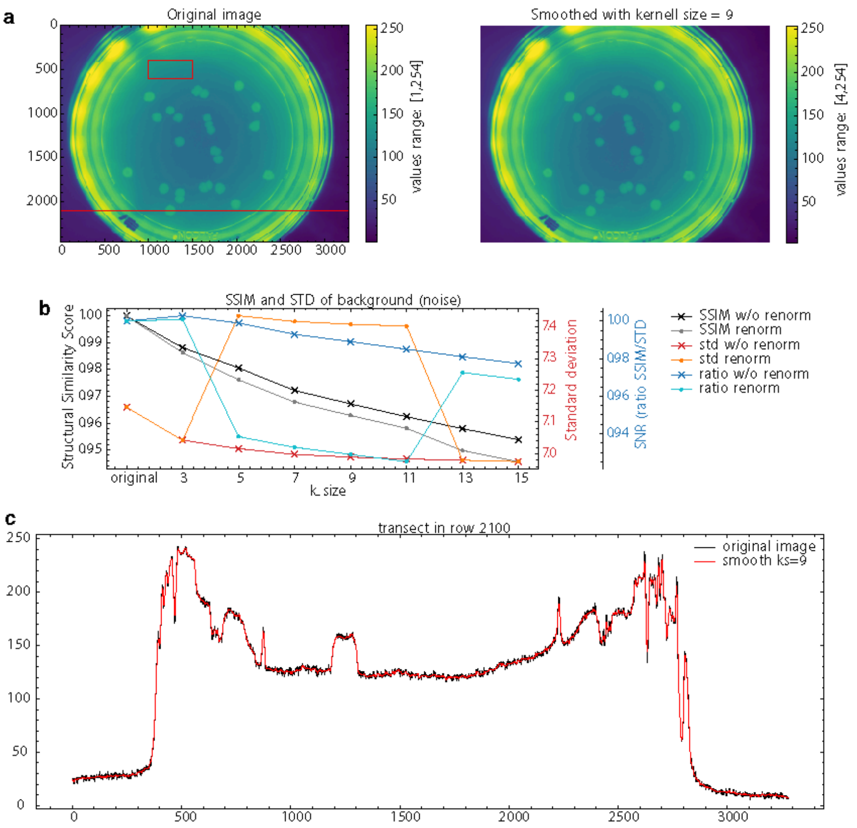


**Figure S8: Interactive filtering procedure. a**) Comparison between the original raw image (left) and the Gaussian filtered image (right), with the corresponding kernel size selected by the user. **b)** Structural Similarity and Standard Deviation (SSIM and STD) of the background, used as a proxy to quantify the information lost by smoothing. **c)** Transect of the selected row of the image, comparing the original values and the filtered values.

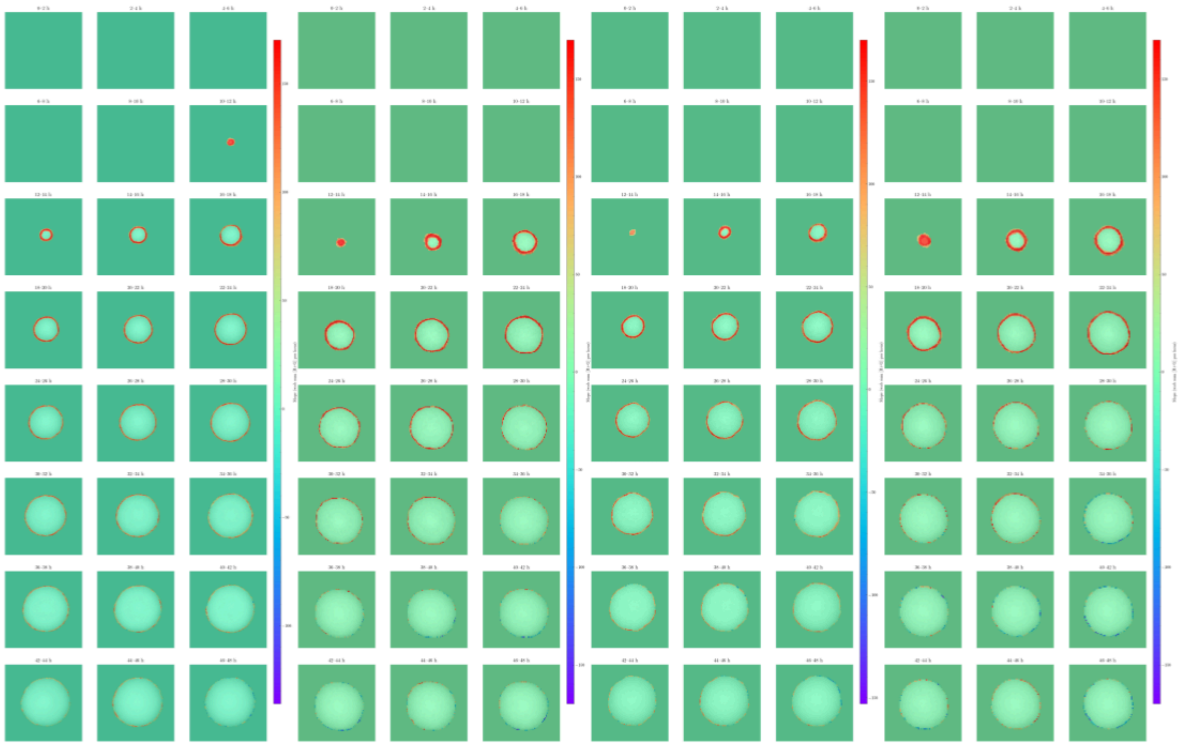

**Figure S9: Rates of GFP expression under constant green light stimulation**. Rates were calculated using 2 hour blocks (8 data points per block) by taking the difference between the fluorescence images of the ROI at each point and averaging them across the block. Heatmap: fluorescence as pixel intensity per hour.

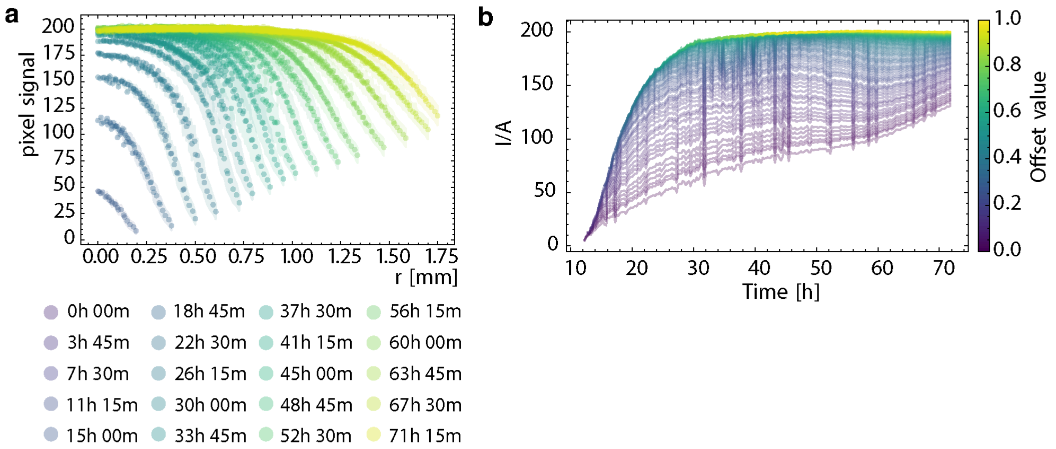


**Figure S10: Spatial distribution of optogenetic responses in colonies induced with green light constantly for 72h.** a) Fluorescence intensity across the colony at different time points. b) Intensity per area per time at different positions from the colony edge (offset), from 0 to 1 mm.

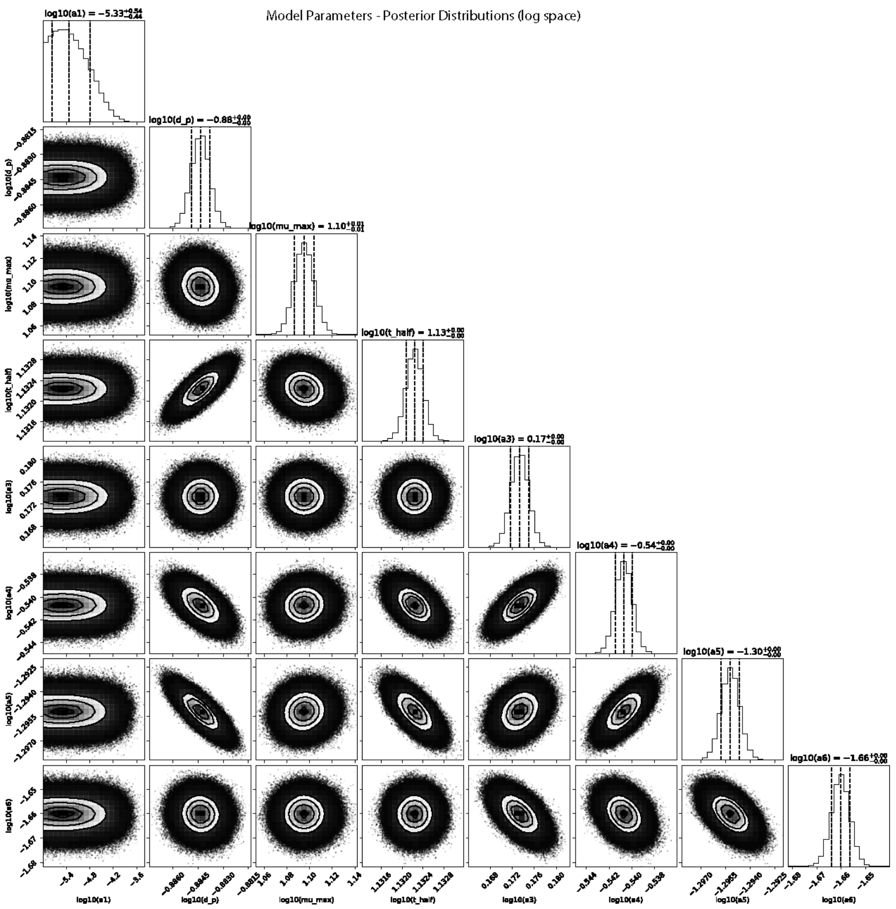


**Figures S11: Posterior parameter distributions for the ODE model (log-space)**. Corner plot summarizing the marginal and pairwise posterior distributions for the parameters of the optogenetic ODE model. Diagonal panels correspond to the 1D posterior distribution histograms, with dashed vertical lines indicating the 16th, 50th and 84th percentiles. Each histogram reports the median and the credible interval. Parameters are represented in base -10 logarithmic scale as the sampling was enforced in log-10 space.

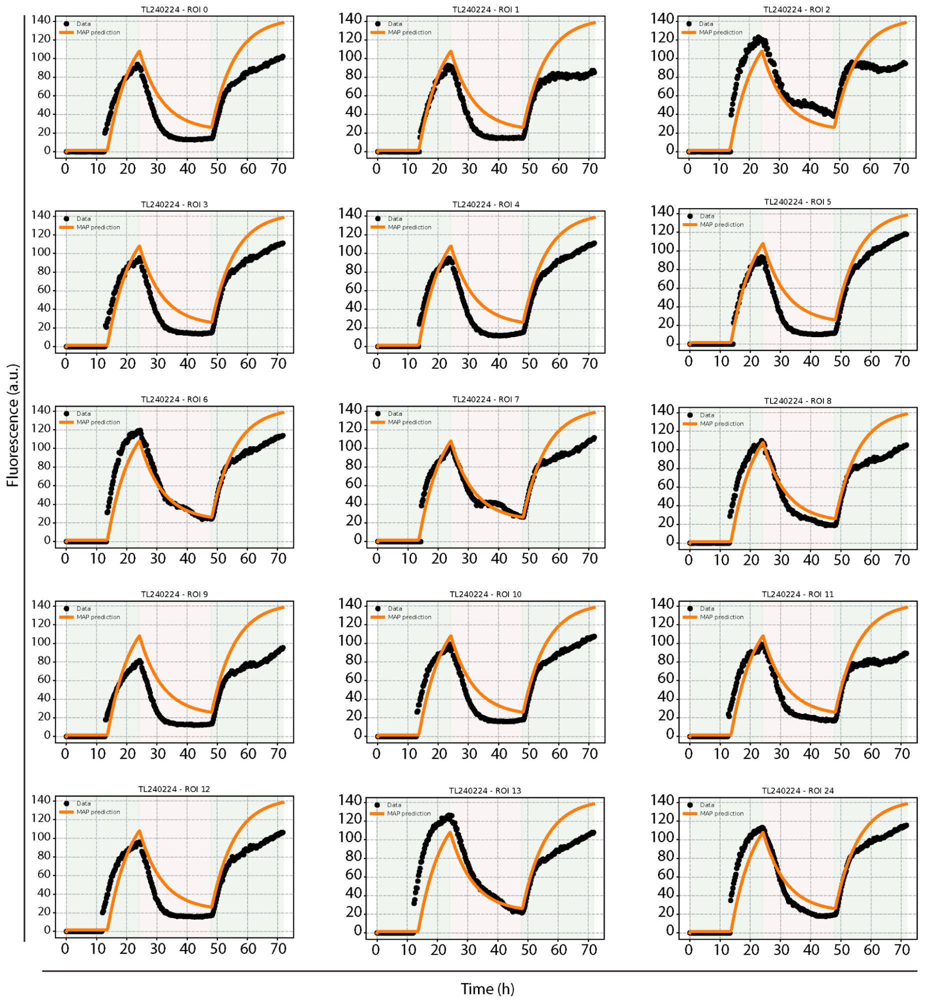


**Figure S12: Model fitting of all colonies in a petri dish.** Representative plots showing experimental data (black) (calculated with width=5px;offset=1px) and model prediction (orange) using the estimated parameters. Data corresponds to the TL240224 dataset, with an optogenetic regime of 0-24 hours of green light, 24-48 hours of red light, and 48-72 hours of green light (shaded regions in each plot).

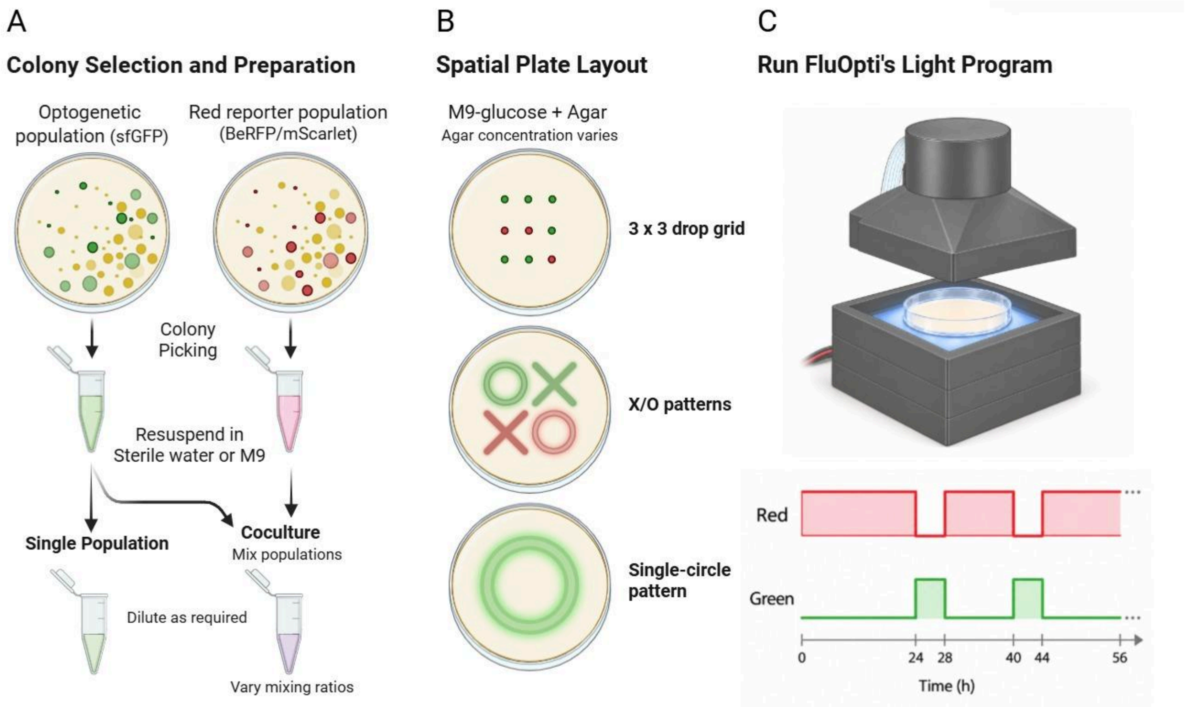


**Figure S13. General pipeline protocol for FluOpti related essays, single or cocultures.** (A) Colony selection and preparation. (B) Spatial inoculation as a 3 x 3 drop grid, X/O patterns, or a single circle. (C) FluOpti operation with an example red/green light program, shaded areas indicate active illumination. Figure created with BioRender.com.

### Supplementary Table

| Experiment ID | Light regime | Colonies (n) | Exponent β [95% CI] | $R^2$ (log10) |
|---|---|---|---|---|
| TL020224 | G 100%, 20 h then R 100%, 28 h | 28 | 2.783 [2.098, 3.467] | 0.729 |
| TL240227 | G 100%, 72 h | 32 | 0.708 [-0.040, 1.457] | 0.111 |
| TL020324 | R 100%, 36 h then G 100%, 36 h | 43 | 3.062 [2.598, 3.527] | 0.812 |
| TL050324 | G 100%, 24 h then R 100%, 48 h | 27 | 1.300 [0.974, 1.625] | 0.730 |
| TL080324 | R 100%, 72 h | 20 | 3.260 [1.636, 4.884] | 0.497 |
| TL240521 | G 20%, 48 h | 467 | 1.390 [1.240, 1.540] | 0.416 |
| TL240523 | G 40%, 48 h | 111 | 1.025 [0.618, 1.432] | 0.186 |
| TL240526 | G 60%, 48 h | 323 | 1.174 [1.001, 1.347] | 0.357 |
| TL240528 | G 80%, 48 h | 66 | 1.676 [0.838, 2.513] | 0.200 |
| TL240530 | Dark | 106 | 2.681 [2.220, 3.141] | 0.561 |
| TL240601 | R 100%, approximately 60 h | 236 | 1.693 [1.551, 1.835] | 0.703 |
| TL240607 | R 100% + G 100%, 72 h | 276 | 1.204 [1.108, 1.300] | 0.690 |

| | | | | |
|---|---|---|---|---|
| TL240610 | R 50%, 48 h | 154 | 1.519 [1.231, 1.806] | 0.418 |
| TL240615 | R 75%, 48 h | 250 | 1.560 [1.397, 1.723] | 0.589 |
| TL240627 | G 100%, 72 h | 256 | 1.618 [1.505, 1.731] | 0.757 |
| TL240716 | R 10% + G 100%, 48 h | 93 | 1.123 [0.577, 1.669] | 0.155 |
| TL240719 | R 10% + G 50%, 48 h | 54 | 3.035 [1.849, 4.222] | 0.336 |
| TL240727 | R 10% + G 25%, 48 h | 203 | 2.110 [1.864, 2.355] | 0.588 |
| TL240730 | R 50% + G 50%, 48 h | 102 | 2.419 [2.023, 2.815] | 0.595 |
| TL240801 | R 20% + G 50%, 48 h | 170 | 2.008 [1.697, 2.319] | 0.491 |
| TL240804 | R 30% + G 50%, 48 h | 125 | 2.256 [1.919, 2.593] | 0.588 |
| TL240806 | R 40% + G 50%, 48 h | 210 | 1.955 [1.692, 2.218] | 0.508 |
| **Pooled** | **All 22 datasets** | **3,352** | **1.390 [1.349, 1.431]** | **0.568** |

**Table S1:** Voronoi segmentation of agar territory available to each colony. Number of cells per plates vs Voronoi cell area and colony size. (22 datasets, 3.352 colonies) Ordinary Linear Squares (OLS) Model: $\log10(\text{Voronoi area}) = \alpha + \beta \log10(\text{colony area})$; areas in $px^2$. 95% CIs describe within-dataset OLS (Ordinary Linear Squares) fits. $R^2$ is calculated in log10.

### Supplementary Videos

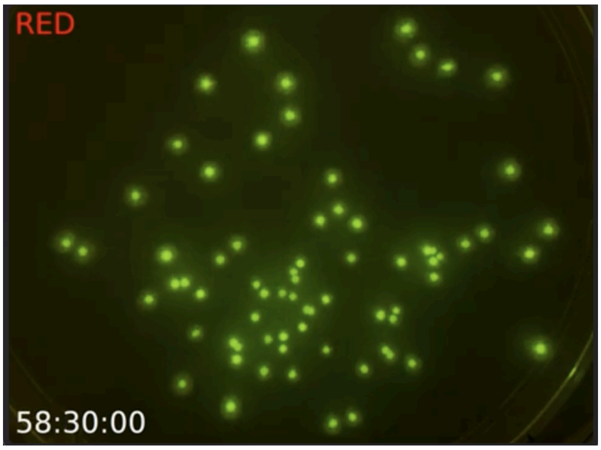


**S1 Video: Time lapse of whole plate fluorescence imaging** https://youtu.be/ohrAiSfcnf4

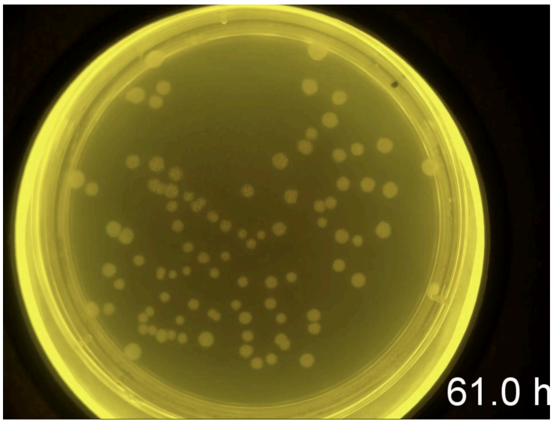


**S2 Video: Time lapse of whole plate brightfield imaging**: https://youtu.be/vb18zDGv9pQ

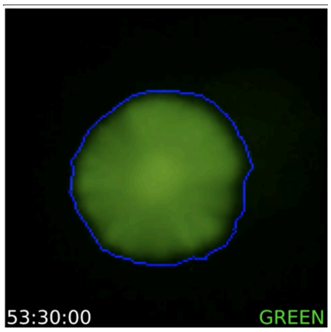


**Video S3: Colony growth computed by edge detection algorithms.**
https://youtube.com/shorts/L93wIbHWHn8?feature=share

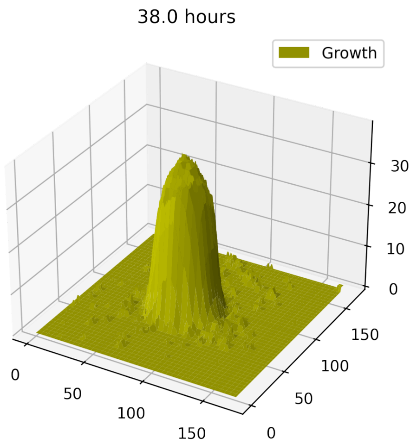


**Video S4:Colony growth.** https://www.youtube.com/shorts/yUkCqt46HQQ

### Supplementary note 1

Jupyter notebook containing the full image processing and analysis pipeline. This notebook is available as a Google Colab in the link (https://colab.research.google.com/github/LabTecLibres/FluOpti/blob/master/data_processing/01_RawDatasetProcessing.ipynb), and also available at the FluOpti repo here: https://github.com/LabTecLibres/FluOpti/tree/master/data_processing

**Supplementary note 2**

Text describing the full derivation of the mathematical model for predicting light responses by the CcaR/CcaS system. Also contains the parameter estimation methods, posterior distributions and descriptions of each parameter and their biological relevance. Supplementary_Note_2_FluOpti_Math_Model.pdf. Code for simulating the model and parameter estimation can be found at the FluOpti repo here: https://github.com/LabTecLibres/FluOpti/tree/master/data_processing
